# Retinal vibrations in bacteriorhodopsin are mechanically harmonic but electronically anharmonic: evidence from overtone and combination bands

**DOI:** 10.1101/2021.07.28.454158

**Authors:** Victor A. Lorenz-Fonfria, Kiyoshi Yagi, Shota Ito, Hideki Kandori

## Abstract

Vibrations of the chromophore in the membrane protein bacteriorhodopsin (BR), a protonated Schiff base retinal, have been studied for decades, both by resonance Raman and by infrared (IR) difference spectroscopy. In spite the light-induced IR difference spectrum between the K intermediate (13-cis retinal) and the initial BR state (all-trans retinal) being first published almost 40 years ago, we present here unreported bands in the 2500 to 1800 cm^−1^ region. We show that the bands between 2500 and 2300 cm^−1^ originate from overtone and combination transitions of retinal C-C stretches. We assigned some of the newly reported bands below 2300 cm^−1^ to the combination of retinal C-C stretches with methyl rocks and with hydrogen-out-of-plane vibrations. Remarkably, experimental C-C overtone bands appeared at roughly twice the wavenumber of their fundamentals, with anharmonic mechanical constants ≤ 3.5 cm^−1^, and in some cases of ∼1 cm^−1^. Comparison of combination and fundamental bands indicates that most of the mechanical coupling constants are also very small. Despite the mechanical quasi-harmonicity of the C-C stretches, the area of their overtone bands was only ∼50 to ∼100 times smaller than of their fundamental bands. We concluded that electronic anharmonicity, the second mechanism giving intensity to overtone bands, must be particularly high for the retinal C-C stretches. We corroborated the assignments of negative bands in the K-BR difference spectrum by ab initio anharmonic spectral calculations of all-trans retinal in BR, which also reproduced reasonably well the small experimental anharmonic and coupling mechanical constants. Yet, and in spite accounting for both mechanical and electronic anharmonicities, the intensity of overtone C-C transitions was underestimated by a factor of 4 to 20, indicating room for improvement in state-of-the-art anharmonic vibrational calculations.

## 1 Introduction

Infrared (IR) spectroscopy is based on the interaction of the electromagnetic field with quantized vibrational levels of molecules. These energy levels, and their associated wavefunctions, dependent on the shape of the multidimensional potential energy surface holding the atoms of the molecule together (Struve, 1989). Because of its deep connection with chemical bonds, IR spectroscopy holds a prominent position in the characterization of molecules, from classical studies on diatomic molecules in the gas phase(Struve, 1989), to more modern applications on biological macromolecules (Koziol et al., 2015; Ghosh et al., 2017). Particularly successful has been the application of IR difference spectroscopy to light-sensitive proteins, where spectral changes following light excitation are selectively resolved, often as a function of time (Kötting and Gerwert, 2013; Kottke et al., 2017). Nevertheless, the interpretation of changes in positions, intensities and/or widths of bands in IR difference spectra of proteins is often based on simple rules of thumb about the effects of H-bonding, polarity and vibrational coupling on vibrational properties (Barth and Zscherp, 2002; Lorenz-Fonfria, 2020), rarely specific enough to provide quantitative atomistic predictions. In this context, vibrational calculations are gaining popularity as a tool to guide the interpretation of experimental spectra in atomic terms, resolving ambiguities about protonation states and H-bonds of groups (Domratcheva et al., 2016; Nakamura and Noguchi, 2016, 2017; Peuker et al., 2016), or tautomers (Domratcheva et al., 2016), complementing ambiguous or missing information in X-ray crystallographic structures of proteins. However, the utility of vibrational computations to interpret experimental IR spectra from biomolecules is still limited by our ability to accurately calculate IR spectra from input molecular structures.

One common approximation in vibrational calculations is the assumption that atomic displacements from the energy minimum in a vibrational mode are small and, thus, that the potential energy surface can be expanded along selected molecular coordinates (often the normal mode coordinates) discarding higher than quadratic terms. This approximation, known as mechanical harmonicity, leads to harmonic wavefunctions. In addition, the calculation of the IR intensities, related to the probability of a transition, often assumes that the dipole moment of the molecule changes linearly along the selected coordinate, an approximation known as electronic harmonicity. These two approximations, known as the harmonic oscillator linear dipole approximation, leads to the familiar quantum harmonic oscillator (Struve, 1989), where only fundamental transitions (0 ⟶ 1) and hot transitions (*i* ⟶ *i* ± 1, with *i* > 0), the latter rare at room temperature, are allowed by selection rules. An experimental hallmark for the breakdown of the harmonic approximation is the observation of overtone bands, *i*.*e*., transitions from the vibrational ground state to the second and further excited states (0 ⟶ *n*, with *n* = 2, 3…). Another hallmark are combination bands, *i*.*e*., light absorption simultaneously driving two vibrations from their vibrational ground state to the first ([0,0] ⟶ [1,1]) or further excited states ([0,0] ⟶ [1,2]; [0,0] ⟶ [2,1]; [0,0] ⟶ [2,2], etc.).

Mechanical anharmonicity can be easily diagnosed and quantified because it affects the frequency difference between fundamental and overtone transitions. For instance, the mechanical anharmonic constant of mode *i, X*_*ii*_, can be determined as the fundamental minus half the first overtone frequency (Sandorfy et al., 2006). Mechanical anharmonicities have been most studied for X-H stretching vibrations (X = C, N, O, S), typically ranging from 50 to 85 cm^−1^ when free from H-bonding (Struve, 1989). Data for other vibrations is scarcer. For X-Y stretches (X and Y = C, N or O) of diatomic molecules (Struve, 1989) or for C=O stretches from carbonyls (Groh, 1988), mechanical anharmonic constants are considerably smaller, between 12-14 cm^−1^, and for C-X stretches (X = F or Cl), as small as 5 cm^−1^ (Groh, 1988). On the other hand, the mechanical coupling constant between the *i* and *j* modes, *X*_*ij*_, is the sum of the *i* and *j* fundamentals minus the frequency of their first combination band (Sandorfy et al., 2006). As a reference, the coupling constant between symmetric and asymmetric X-H vibrations (X= N or O) is around 90-160 cm^−1^ (Sandorfy et al., 2006).

Mechanical anharmonicity not only modifies the energy of vibrational levels and, thus, the frequency of fundamental and overtone/combination transitions, but it also modifies the wavefunction of vibrational states. Consequently, it gives IR intensity (absorption probability) to overtone and combination transitions, forbidden (i.e., with null IR intensity) under the harmonic approximation. However, while the presence of overtone and combination bands is often considered a sign of mechanical anharmonicity, electronic anharmonicity alone is sufficient to give IR intensity to overtones and combination bands, even for purely harmonic wavefunctions, affecting even the intensity of fundamental transitions (Sandorfy et al., 2006; Panek and Jacob, 2016). Because the IR intensity of overtone and combination bands is simultaneously affected by mechanical and electronic anharmonicities, in practice both contributions are experimentally difficult to disentangle from each other. Consequently, the role played by electronic anharmonicities on shaping IR spectra in biological macromolecules is yet to be demonstrated experimentally to our knowledge.

Bacteriorhodopsin (BR) is a light-driven proton-pump membrane protein that belongs to the family of microbial rhodopsin, a family present in the genome of cells in all kingdoms of life (Ernst et al., 2014). It contains seven transmembrane helices and a protonated Schiff base (PSB) retinal chromophore (Fig. 1A). Photoisomerization of the PSB retinal from all-trans to 13-cis conformation (Fig. 1B) starts a photocyclic reaction, where a proton is vectorially transported from the cytoplasm to the extracellular side (Ernst et al., 2014). The photocycle of BR involves various intermediate states which can be trapped at low temperatures (Balashov and Ebrey, 2007). The intermediate K, the first thermally metastable state of the photocycle, can be trapped by illumination at 80 K (Rothschild and Marrero, 1982; Balashov and Ebrey, 2007).

**Figure 1.**
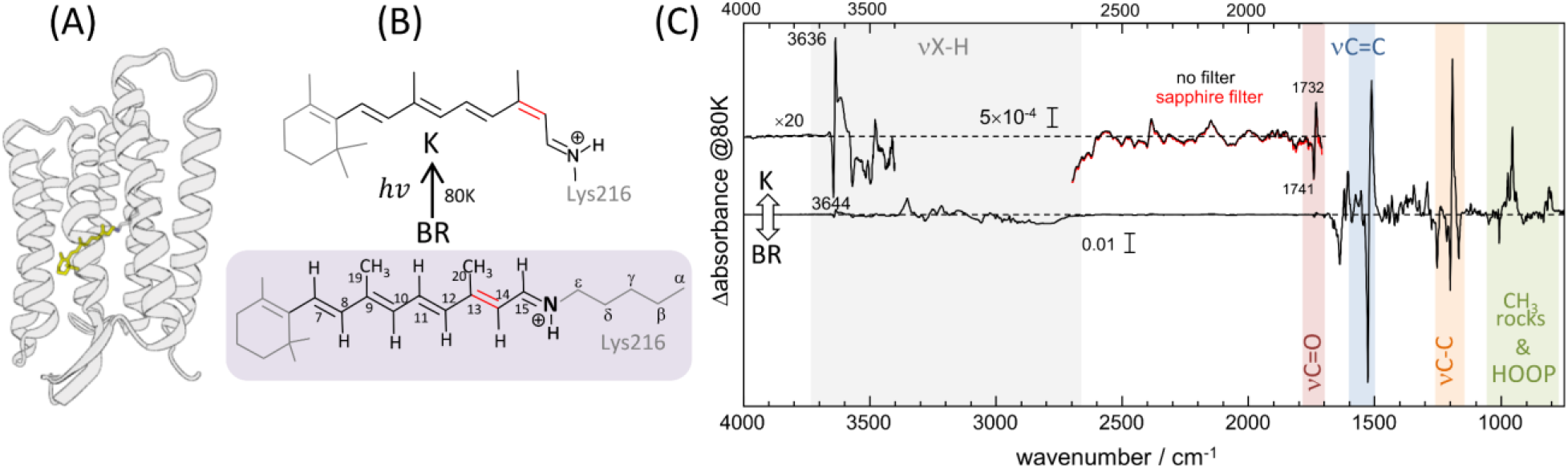
Structural and vibrational differences between bacteriorhodopsin (BR) in its initial state (all-trans retinal) and in the K intermediate (13-cis retinal). (a) Structure of BR in the initial state (pdb 1C3W) (Luecke et al., 1999), highlighting the location of the retinal chromophore (in yellow), forming a protonated Schiff base (PSB) with Lys216. (b) Schematic representation of the PSB retinal chromophore in the initial state of BR (all-trans, 15-anti conformation) and in the K intermediate (13-cis, 15-anti conformation), including the numbering of relevant carbon atoms. (c) K-minus-BR difference FT-IR spectrum obtained by illumination at 80 K, with highlighted regions corresponding to X-H stretches, carboxylic C=O stretches, retinal C=C and C-C stretches, retinal CH_3_ rocks, and retinal hydrogen out of plane (HOOP) ethyl wags. The inset above 3400 cm^−1^, besides illustrating the noise level, shows bands at 3644 (–) and 3636 (+) cm^−1^ from the νO-H of internal dangling water 401. The inset between 2600 and 1710 cm^−1^ shows bands at 1732 (+) and 1741 (–) cm^−1^ from carboxylic νC=O vibrations of Asp115, as well as several bands never reported before. Measurements with a sapphire window, acting as a high pass filter (cutoff at ∼1650 cm^−1^), exclude that double-modulation artifacts might contribute to the absorption changes between 2600 and 1710 cm^−1^.

We report on previously unnoticed bands between 2550-2300 cm^−1^ in the FT-IR difference spectrum between the K intermediate and the initial BR state at 80 K (K-BR for short). Based on their wavenumber, we assigned most of them to overtone and combination bands of C-C stretching vibrations of the retinal: the negative bands to the initial state of BR (all-trans PSB retinal) and the positive bands to the K intermediate (13-cis PSB retinal retinal). Comparing bands from fundamental C-C stretches with their putative overtones, we concluded that retinal C-C stretching vibrations are virtually mechanically harmonic both in BR (all-trans retinal) and in the K intermediate (13-cis retinal), with anharmonic mechanical constants between +1.0 cm^−1^ and +3.5 cm^−1^. The mechanical quasi-harmonicity for the retinal C-C stretches reported here sharply contrasts with the relative area of the overtone bands, only ∼50-100 times smaller than for their fundamentals. We take this observation as indication that retinal vibrations in bacteriorhodopsin are notably electronically anharmonic.

We complemented the experimental observations with two state-of-the-art ab initio anharmonic vibrational calculations for all-trans PSB retinal in the BR initial state. Calculated spectra correctly reproduced experimental bands from fundamental C-C vibrations, as well as many putative overtones/combination bands, helping in their assignment. The calculations also reproduced the small mechanical anharmonic and coupling constants of the C-C stretches. However, they underestimated by a factor of 4-15 the relative intensity of the overtone bands with respect to their fundamentals. This discrepancy might indicate potential limitations in current state-of-the-art anharmonic calculations, for instance when electronic anharmonicities are dominant, as in the present case.

Overall, our results provide a solid example for the role that electronic anharmonicity in determining the intensity of overtone and combination bands. We believe that our experimental results, rich in details, could be useful to benchmark future developments in anharmonic vibrational calculations.

## 2 Materials and Methods

### 2.1. FT-IR difference spectroscopy

Purple membranes were isolated from *Halobacterium salinarum* as described (Oesterhelt and Stoeckenius, 1974). Around 25 μl of a 6 mg/ml solution of BR in purple membranes (3 mM MES, 2 mM NaCl, pH 6.5) was dried at ambient humidity on top of a BaF_2_ window of 18 mm diameter. The resulting film was rehydrated through the vapor phase provided by 4 μl of a mixture of water/glycerol (9/1 w/w) distributed in 3-5 drops placed nearby the film and closed with a second BaF_2_ window using a 0.5 mm thick silicone spacer. The sample was placed on a cryostat (OptistatDN, Oxford) coupled to a FT-IR spectrometer (Cary 670, Agilent) equipped with a MCT detector and purged with dry nitrogen. The BaF_2_ windows were placed roughly perpendicular to the IR beam but slightly tilted. After letting 30-45 min to complete the hydration process, the sample was light-adapted by 2 min of illumination (>530 nm, IR filtered), and cooled down to 80 K. The FT-IR absorption spectrum of the hydrated film of BR at 80 K is shown in Supplementary Figure 1. The light-induced K-BR FT-IR difference spectrum was obtained by 1 minute of illumination using 540 ± 10 nm light. The back-conversion from K to the BR dark-state was done with >670 nm illumination for 1 minute, to obtain the BR-K spectrum. Interferograms at 2 cm^−1^ resolution were collected before and after the illumination to obtain an FT-IR difference spectrum with a total of 128 accumulated scans. Alternating illumination to promote the forward and backward photoconversion was repeated five times, resulting in averaged K-BR and BR-K with 640 coadded scans each (Supplementary Figure 2). Further averaging the K-BR and the minus BR-K difference spectra cancelled trace absorption contributions from water vapor and CO_2_ (Supplementary Figure 2). In spite the high absorbance of the BR film used (Supplementary Figure 1), the obtained K-BR IR difference spectrum was almost indistinguishable from that obtained with a sample with three times less absorbance (Supplementary Figure 3), except at regions with an absorbance higher than 2. When displaying the whole K-BR difference spectrum (Fig. 1c), the absorbance changes in regions with a background absorbance above 2, were taken, after an appropriate scaling, from the sample with three times less protein. K-BR difference spectra were measured with and without a sapphire window placed in the optical path, which fully blocks light below 1600 cm^−1^, without any appreciable discrepancy in the 2600-1800 cm^−1^ region between them. When using an older FT-IR spectrometer (Biorad FTS-40), used in a previous publication (Ito et al., 2018), the use of a sapphire window was required to disentangle true overtone and combination band signals from double modulation artifacts (Supplementary Figure 4).

### 2.2. Anharmonic vibrational calculations

The atomistic model of BR was constructed based on an X-ray crystal structure, with PDB ID 1C3W (Luecke et al., 1999). Missing residues between 157 – 161 were complemented using MODELLER 9.14. (Šali, 2014). The protonation state of titratable residues was determined based on pKa values obtained by PROPKA 3.1. (Olsson et al., 2011): Asp96, Asp115, and Glu194 were protonated, while other residues were kept ionized. Hydrogen atoms were added using the HBUILD utility of CHARMM (Brooks et al., 2009), and relaxed by performing molecular dynamics (MD) simulations for 100 ps with the positions of the heavy atoms kept fixed.

The last structure of the MD trajectory was used for the subsequent QM/MM calculations (Yagi et al., 2019). In the QM/MM calculation, the retinal, as well as three nearby residues (sidechains of Asp85 and Asp212 beyond C_β_ and of Lys216 beyond C_γ_) and a crystal water molecule (Wat402), were included in the QM region (74 atoms in total), treated at the DFT level. For the MM region, which includes the remaining protein and water molecules, we used CHARMM36 (Best et al., 2012) and TIP3P (Jorgensen et al., 1983) force fields, respectively. The energy minimization was followed by harmonic vibrational analysis, with a partial Hessian which includes all atoms from the retinal and from Lys216 beyond C_δ_ (56 atoms, 168 vibrational modes).

Anharmonic vibrational calculations were carried out following two schemes. In the first scheme (scheme A), the vibrational calculation was performed in 116 dimensions neglecting 52 low frequency modes (< 800 cm^−1^). Optimized-coordinate vibrational self-consistent field (oc-VSCF) was first carried out with a cubic force field to optimize the vibrational coordinates (Yagi et al., 2012; Yagi and Otaki, 2014). These coordinates were used to generate the anharmonic potential energy surface (PES). The quartic force field (Yagi et al., 2004) was generated in 116 dimensions. The grid potential (Yagi et al., 2000) was generated in one-dimension for 83 modes with 11 grid points, including C-H and N-H stretching modes, and in two-dimension with 9 grid points for the combination of modes 72/X, 78/X, and X/X, where X represents modes 86, 89, 92, or 93. Note that the dipole moment surfaces (DMS) were obtained simultaneously with the PES. Then, VSCF and the second-order vibrational quasi-degenerate perturbation theory (VQDPT2) calculations (Yagi et al., 2008; Yagi and Otaki, 2014) were performed to obtain an IR line spectrum (transition intensities vs transitions energies). VSCF was carried out with 11 harmonic oscillator basis functions for each mode, and VQDPT2 was carried out setting the maximum sum of excitation to 4 and the number of P space generation to 1. In the second scheme (scheme B), the calculation was performed in only 8 selected modes consisting of four C-C stretching modes (modes 86, 89, 92, and 93), and modes 70, 72, 82, 136. The PES and DMS were constructed using a grid method with 11 and 9 grid points for the one-mode and the two-mode part, respectively (Yagi et al., 2000). Finally, the IR line spectrum was obtained by VSCF and vibrational configuration interaction (VCI) calculations using the resulting PES and DMS. VSCF was performed with 11 harmonic oscillator basis functions on each mode, and VCI was carried out at the level of three-mode excitation, setting the maximum sum of quantum number to 6.

The QM/MM calculations were performed using a development version of GENESIS 1.6.1. (www.r-ccs.riken.jp/labs/cbrt) (Jung et al., 2015; Kobayashi et al., 2017). The QM calculation was carried out at the level of B3LYP-D3 (Lee et al., 1988; Becke, 1993; Grimme et al., 2010) with cc-pVDZ (scheme A) or cc-pVTZ (scheme B) basis sets (Dunning, 1989) using Gaussian16 (http://gaussian.com/gaussian16/) (Frisch et al., 2016) or Q-Simulate (see https://qsimulate.com/), as done previously (Yagi et al., 2021). The vibrational calculation was performed using SINDO 4.0 (available at tms.riken.jp/en/research/software/sindo).

### 2.3. Data postprocessing

IR line spectra, obtained from anharmonic vibrational calculations, were convoluted with a Lorentzian band to facilitate their comparison with the experimental spectral. For fundamental transitions we used a Lorentzian full width at half height (FWHH) of 7 cm^−1^, and for overtone and combination transitions a FWHH of 14 cm^−1^. This choice reflects the experimental observation that overtone bands appear twice broader than fundamental bands. On the other hand, to improve the resolution of overlapping bands, the experimental K-BR difference FT-IR spectrum was band-narrowed by Fourier self-deconvolution (Kauppinen et al., 1981) as implemented in the Matlab App FourierDataProcessing, available at https://www.mathworks.com/matlabcentral/fileexchange/92573-fourierdataprocessing.

## 3 Results and Discussion

### 3.1. K-minus-BR FT-IR difference spectrum

Light-induced FT-IR difference spectra of BR at low temperature have been amply measured and characterized in the past (Bagley et al., 1982; Rothschild and Marrero, 1982; Siebert and Mäntele, 1983; Earnest et al., 1986; Fischer et al., 1994; Kandori et al., 1998), as reviewed (Maeda, 1995; Kandori, 2020). Illumination at 80 K with 540 nm traps the K intermediate (Rothschild and Marrero, 1982; Balashov and Ebrey, 2007), the first metastable intermediate after all-trans to 13-cis photoisomerization of the PSB retinal. The trapped K intermediate reverts to the BR state with >670 nm illumination (Rothschild and Marrero, 1982; Balashov and Ebrey, 2007). Figure 1c shows the K-minus-BR FT-IR difference spectrum (K-BR spectrum for short), where positive bands correspond to the K intermediate (13-cis PSB retinal) and negative bands correspond to the initial BR state (all-trans PSB retinal), BR state for short. In the K intermediate most of the structural changes are restricted to the chromophore and its vicinity (Neutze et al., 2002; Wickstrand et al., 2019), and, thus, most of the resolved bands in the K-BR spectrum come from vibrational modes located at the retinal molecule (Rothschild et al., 1984). At ∼1600-1500 cm^−1^ and at ∼1275-1150 cm^−1^ we find bands from fundamental transitions of retinal C=C and C-C stretches, and between 1050 and 750 cm^−1^ fundamental transitions of methyl rocks and hydrogen out of plane (HOOP) vibrations (Smith et al., 1987; Maeda, 1995). The specific assignment of bands between 1275 and 900 cm^−1^ is given in Fig. 2 (blue trace), and assignments for the negative bands are collected in Table 1. Besides retinal vibrations, we have bands between 1780-1700 cm^−1^ from fundamental transitions of the carboxylic C=O stretching of Asp115 (Braiman et al., 1988), and between ∼3700 and ∼2700 cm^−1^ from fundamental transitions of X-H vibrations (X = O, N, and C), including those above 3650 cm^−1^ from the O-H stretch of the dangling internal water 401 (Hayashi et al., 2004).

**Figure 2.**
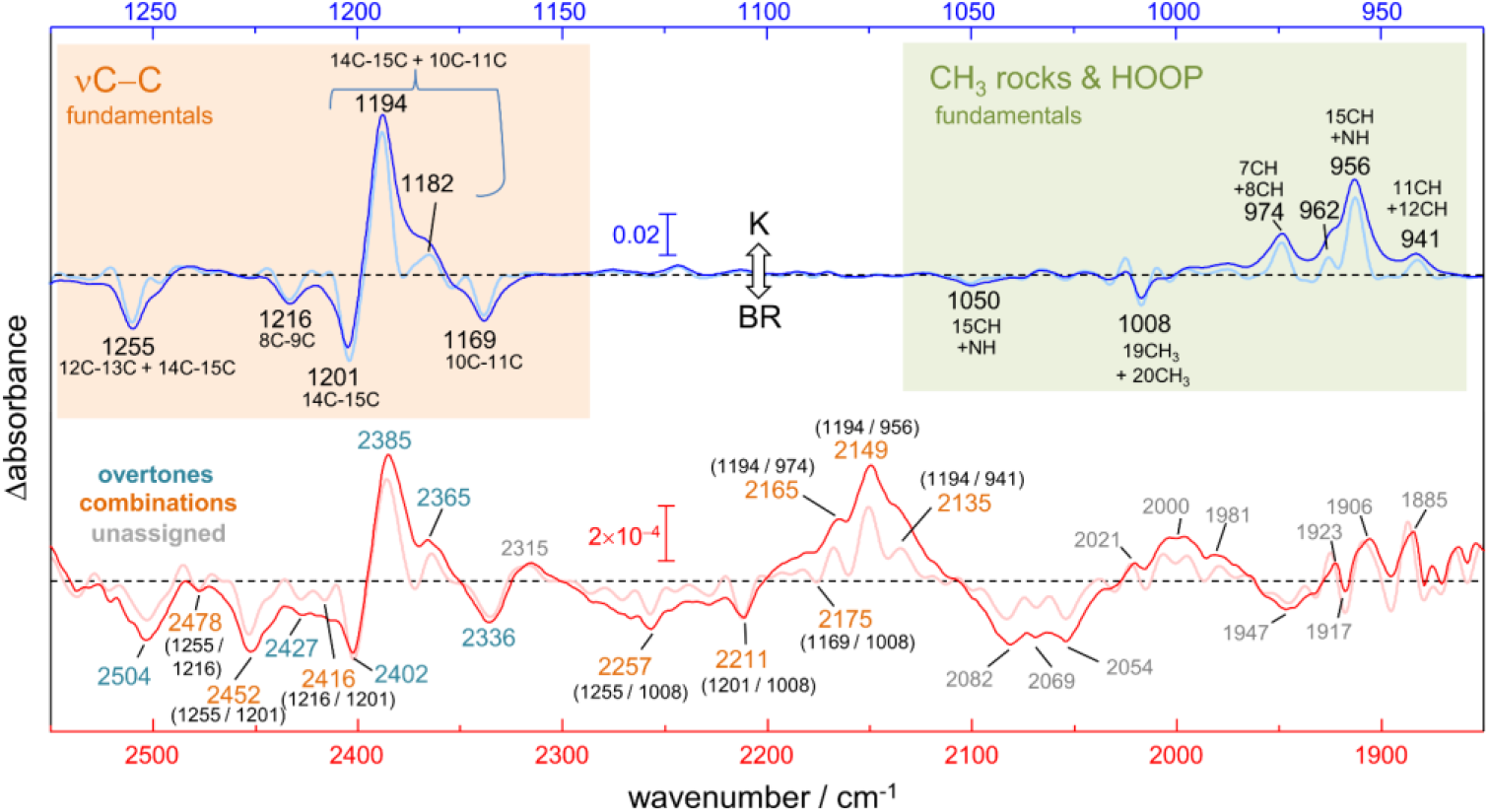
K-BR difference FT-IR spectrum at 80 K. Positive bands come from vibrations in the K intermediate and negative bands in the initial BR state. The figure compares fundamental transitions from retinal C-C stretches, CH_3_ rocks and HOOP vibrations in the 1275-925 cm^−1^ region (blue trace), with potential overtones and combinations in the 2550-1850 cm^−1^ region (red trace). The K-BR spectrum is also displayed after band narrowing with Fourier self-deconvolution (pale blue and pale red traces). For combination bands, the wavenumber of the two putative fundamental frequencies is given in brackets.

**Table 1.** Assignment of bands to fundamental vibrational transitions of the retinal in the BR state.

| assignment | experiment |  | scheme A<sup>c</sup> |  |  | scheme B<sup>d</sup> |  |  |
| --- | --- | --- | --- | --- | --- | --- | --- | --- |
| | $\nu$ / cm<sup>-1</sup> | Area (relative)<sup>a</sup> | State | $\nu$ / cm<sup>-1</sup> ($\Delta\nu$)<sup>e</sup> | Intensity (relative)<sup>b</sup> | State | $\nu$ / cm<sup>-1</sup> ($\Delta\nu$) | Intensity (relative) |
| 19C/20C rock | 1008 | 0.06 (0.20) | 72<sub>1</sub> | 1002 (-6) | 191 (0.49) | 72<sub>1</sub> | 1034 (+26) | 123 (0.36) |
| H15C=NH<br>HOOP | 1050 | 0.09 (0.30) | 82<sub>1</sub> | 1040 (-10) | 73 (0.19) | 82<sub>1</sub> | 1108 (+58) | 64 (0.18) |
| 10C-11C | 1169 | 0.29 (1.0) | 86<sub>1</sub> | 1199 (+30) | 389 (1.0) | 86<sub>1</sub> | 1211 (+42) | 345 (1.0) |
| 14C-15C | 1201 | 0.59 (2.0) | 89<sub>1</sub> | 1232 (+31) | 888 (2.3) | 89<sub>1</sub> | 1252 (+51) | 261 (0.76) |
| 8C-9C | 1216 | 0.18 (0.62) | 92<sub>1</sub> | 1225 (+9) | 28 (0.07) | 92<sub>1</sub> | 1264 (+48) | 771 (2.2) |
| 12C-13C / 14C-15C | 1255 | 0.37 (1.2) | 93<sub>1</sub> | 1269 (+14) | 649 (1.7) | 93<sub>1</sub> | 1300 (+45) | 297 (0.86) |

<sup>a</sup> Experimental area in the K-BR spectrum, in cm<sup>-1</sup> (in brackets relative to the 10C-11C fundamental
<sup>b</sup> Calculated intensity km/mol (in brackets relative to the intensity of the state 86<sub>1</sub>)
<sup>c</sup> VQDPT2@B3LYP-D3/cc-pVDZ
<sup>d</sup> VCI@B3LYP-D3/cc-pVTZ
<sup>e</sup> $\Delta\nu$ , variation between the experimental and calculated wavenumber

### 3.2. Bands in the 2550-1800 cm^−1^ spectral range

In the K-BR difference spectrum we find previously unreported bands, extending from 2550 to 1800 cm^−1^ (Fig. 1C, insert), in a region where bands from fundamental vibrational transitions rarely appear in proteins (Adhikary et al., 2017). These bands are reproduced in both in the K-BR and in the BR-K difference spectrum (Supplementary Figure 2) and display an intensity far above the noise level (Fig. 1C, Insert). In addition, they are unaffected by placing in the optical path a sapphire window acting as a high pass optical filter (Fig. 1C, insert, compare red and black traces), discarding that they could be ghost bands originated from double modulation artifacts in the interferometer (Ito et al., 2018).

Although fundamental transitions from stretches of triple bonds (e.g., C≡C) and S-H groups can contribute to this spectral region, such chemical groups are absent in BR. We also expect contributions from fundamental transitions of X-D stretches in this spectral range. However, potential bands from X-D stretches are expected to be at least 100 times smaller than the bands observed here upon considering the natural abundance of deuterium (0.016%) and the intensity of X-H bands in the K-BR difference spectrum (Fig. 1C).

Upon reasonably discarding that these new bands could originate from fundamental transitions, we explored their assignment to overtone and/or combination transitions. For that, we compared the spectral features between 2550 and 1800 cm^−1^ (Fig. 2, red trace) with those between 1275 and 900 cm^−1^ (Fig. 2, blue trace). To further facilitate the spectral comparison of both regions, we mathematically narrowed the K-BR spectrum using Fourier self-deconvolution, FSD (Fig. 2, pale blue and pale red traces). Laying both spectral regions on top of each reveals some remarkable similitudes between them (Fig. 2). Several bands that align nicely with each other can be reasonably assigned, based on their wavenumber and appearance, to overtones of retinal C-C stretches (Fig. 2, bottom, cyan labels). On a similar basis, several bands can be reasonably assigned to the combination of fundamentals, either between retinal C-C stretches, between retinal C-C stretches and methyl rocks, or between retinal C-C stretches and HOOP vibrations (Fig. 2, bottom, orange labels). These assignments are collected in Table 2 for the BR state (negative bands), being further developed and justified in view of anharmonic spectral calculations presented in forthcoming sections. But before that, we will briefly review in the next section the assignment of bands in the 1275 and 900 cm^−1^ region to fundamental transitions from vibrations of the PSB retinal.

**Table 2.** Assignment of bands to overtone and combination vibrational transitions of the retinal in the BR state.

| assignment | experiment |  |  |  | State | scheme A |  |  |  | State | scheme B |  |  |  |
| --- | --- | --- | --- | --- | --- | --- | --- | --- | --- | --- | --- | --- | --- | --- |
| | $\nu / \text{cm}^{-1}$ | $X^a / \text{cm}^{-1}$ | Area<sup>d</sup> | F/O<sup>b</sup> | | $\nu / \text{cm}^{-1}$ | $X^a / \text{cm}^{-1}$ | I<sup>d</sup> | F/O<sup>b</sup> | | $\nu / \text{cm}^{-1}$ | $X^a / \text{cm}^{-1}$ | I<sup>d</sup> | F/O<sup>b</sup> |
| 19C/20C rock + 10C-11C | 2175 | +1.7 | 0.0009 |  | 72;86<sub>1</sub> | 2201 (+26) | -0.1 | 0.25 |  | 72;86<sub>1</sub> | 2248 (+73) | -3.2 | 0.21 |  |
| 19C/20C rock + 14C-15C | 2211 | -1.7 | 0.0036 |  | 72;89<sub>1</sub> | 2235 (+24) | +0.1 | 0.88 |  | 72;92<sub>1</sub> | 2302 (+91) | -3.8 | 1.29 |  |
| 19C/20C rock + 12C-13C/14C-15C | 2257 | +6.4 | 0.0020 |  | 72;93<sub>1</sub> | 2271 (+13) | +0.3 | 0.19 |  | 72;93<sub>1</sub> | 2337 (+80) | -2.8 | 0.17 |  |
| (10C-11C)<sub>2</sub> | 2336 | +1.5 | 0.0059 | 50 | 86<sub>2</sub> | 2395 (+59) | +0.9 | 1.53 | 260 | 86<sub>2</sub> | 2422 (+86) | -0.2 | 1.43 | 240 |
| 10C-11C + 14C-15C | ND<sup>c</sup> | - |  |  | 89;86<sub>1</sub> | 2430 | +0.6 | 0.78 |  | 89;86<sub>1</sub> | 2466 | -3.5 | 0.48 |  |
| 10C-11C + 8C-9C | ND | - |  |  | 92;86<sub>1</sub> | 2423 | +1.3 | 0.08 |  | 92;86<sub>1</sub> | 2478 | -3.2 | 1.23 |  |
| (14C-15C)<sub>2</sub> | 2402 | +1.0 | 0.0073 | 80 | 89<sub>2</sub> | 2456 (+54) | +3.9 | 0.50 | 1800 | 89<sub>2</sub> | 2503 (+101) | -0.1 | 0.79 | 330 |
| 8C-9C + 14C-15C | 2416 | +2.7 | 0.0015 |  | 92;89<sub>1</sub> | 2462 (+56) | -5.1 | 1.11 |  | 92;89<sub>1</sub> | 2519 (+103) | -3.1 | 0.47 |  |
| (8C-9C)<sub>2</sub> | 2427 | +2.8 | 0.0016 | 110 | 92<sub>2</sub> | 2450 (+23) | +0.6 | 0.08 | 340 | 92<sub>2</sub> | 2529 (+102) | -0.5 | 0.69 | 1700 |
| 14C-15C + 12C-13C/14C-15C | 2452 | +3.9 | 0.0068 |  | 93;89<sub>1</sub> | 2498 (+46) | +2.1 | 0.92 |  | 93;89<sub>1</sub> | 2556 (+104) | -4.4 | 0.52 |  |
| 8C-9C + 12C-13C/14C-15C | 2478 | +6.1 | 0.0010 |  | 93;92<sub>1</sub> |  | +1.2 | 0.02 |  | 93;92<sub>1</sub> | 2567 (+89) | -3.3 | 0.70 |  |
| (12C-13C/14C-15C)<sub>2</sub> | 2504 | +3.5 | 0.0064 | 60 | 93<sub>2</sub> | 2535 (+31) | +1.1 | 0.67 | 970 | 93<sub>2</sub> | 2604 (+100) | -1.8 | 0.70 | 420 |

<sup>a</sup> X represents the mechanical anharmonic constant for overtones, and the coupling constant for combinations.
<sup>b</sup> Fundamental to overtone area ratio (for experimental spectra) and intensity ratio (for calculated spectra).
<sup>c</sup> ND: not detected
<sup>d</sup> Band area in $\text{cm}^{-1}$ , and IR intensity in ( $\text{km}/\text{mol}$ )
<sup>e</sup> $\Delta\nu$ , variation between the experimental and calculated wavenumber

### 3.3. Fundamental transitions from vibrations of the PSB retinal

We start our assignment with the retinal C-C stretches. For the assignment of the negative bands in Fig. 2 (blue spectrum), we relayed on previous works using BR reconstituted with isotopically labelled retinal (Gerwert and Siebert, 1986; Smith et al., 1987) and vibrational harmonic calculations of the PSB retinal in the BR state (Smith et al., 1987; Babitzki et al., 2009). The bands at 1201 cm^−1^ and at 1169 cm^−1^ correspond mostly to a 14C-15C stretching and a 10C-11C stretching vibration of the PSB retinal, respectively (Gerwert and Siebert, 1986; Smith et al., 1987). You can consult in Fig. 1B the numbering of atoms in the retinal. The negative band at 1255 cm^−1^ has been assigned to coupled 12C-13C and 14C-15C stretches, with additional contributions from in-plane methyl rocks of 14C and 15C and to in-plane methylene vibrations of C_ε_ of Lys216 (Smith et al., 1987; Babitzki et al., 2009). Finally, the small negative band at 1216 cm^−1^ corresponds mostly to a 8C-9C stretch, with some 14C-15C and 9C-19C character (Smith et al., 1987). The negative band at 1008 cm^−1^ (Fig. 2, blue spectrum) has been assigned to a symmetric in-plane rocking combination of the 19C and 20C methyl groups of all-trans PSB retinal (Smith et al., 1987; Yamazaki et al., 1995). The negative band at 1050 cm^−1^ (Fig. 2, blue spectrum), remains unassigned to our best knowledge. Of our anharmonic vibrational calculations, which we will describe below, assigns a similar band to an in-phase H15C=NH HOOP vibration (see mode 82 in Fig. 3A and in Fig. 3E). In agreement with this tentative assignment, this band vanishes upon deuteration of the NH group of PSB retinal by incubation of BR in D_2_O (Supplementary Figure 5).

**Figure 3.**
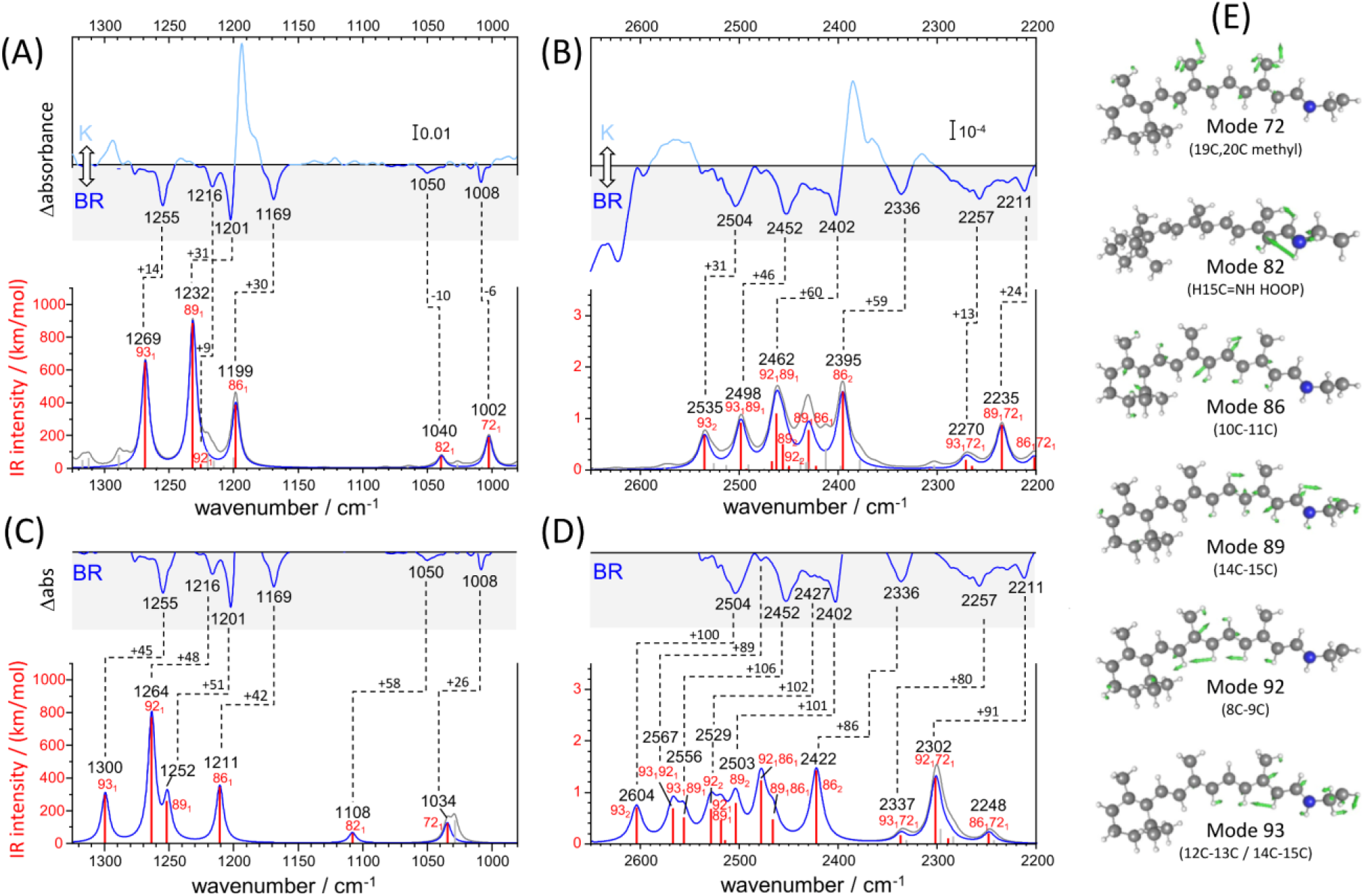
Calculated versus experimental IR spectra of all-trans PSB retinal in BR. (a, b) Calculation with scheme A vs experimental K-BR spectrum. (c, d) Calculation with scheme B vs experimental K-BR spectrum (only the negative bands from the BR state are shown). Bands assigned to a common vibrational transition are connected by dashes lines. See text and Materials and Methods for details about calculations with scheme A and scheme B. (a-d) Wavenumbers and IR intensities of calculated vibrational transitions (gray vertical lines). Transitions from modes with experimentally detected fundamentals (modes 72, 82, 86, 89, 92 and 93) are highlighted in red. Spectra were generated from the calculated intensity of vibrational transitions by convolution with a Lorentzian band (gray trace and blue traces), of either (a,b) 7 cm^−1^ or (c,d) 14 cm^−1^ full width at half height. Transitions are labelled as *x*_1_, *x*_2_, or *x*_1_*y*_1_, for fundamental, overtones and combination transitions, respectively. (e) Atom displacements for some selected vibrational modes.

For the assignment of positive bands, corresponding to 13-cis PSB retinal, we relayed exclusively on previous experimental work using BR reconstituted with isotope labelled retinal (Gerwert and Siebert, 1986; Fahmy et al., 1991; Weidlich and Siebert, 1993), as vibrational calculations for the retinal in the K intermediate are not available to our knowledge. The positive band at 1194 cm^−1^ and the shoulder at 1182 cm^−1^ (Fig. 2, blue spectrum) originates from the mixing of the 14C-15C and 10C-11C stretches of 13-cis PSB retinal (Gerwert and Siebert, 1986). Regarding HOOP bands, at 974, 962, 956 and 941 cm^−1^ in Fig. 2 (blue spectrum), only two of them, at 962 and 941 cm^−1^, are not affected upon deuteration of either N or 15C. Thus, the intense positive band at 956 cm^−1^ has been assigned to the H15C=NH HOOP mode (Maeda et al., 1991; Maeda, 1995). For the remaining bands tentative assignments have been proposed, but with rather weak experimental evidence. For instance, the positive band at the 941 cm^−1^ has been assigned to the H11C=12CH HOOP (Earnest et al., 1986; Weidlich et al., 1997) and the positive band at 974 cm^−1^ to the H7C=8CH HOOP mode (Fahmy et al., 1991).

### 3.4. Anharmonic vibrational calculations of all-trans PSB retinal

We conducted two anharmonic vibrational calculations of all-trans PSB retinal in the BR state, following two schemes: scheme A (Fig. 3A-B) and scheme B (Fig. 3C-D). The technical characteristics of these two anharmonic calculations are detailed in Materials and Methods. Briefly, our calculations started with a QM/MM energy minimization of the BR structure, with the QM region including the retinal, Lys216 from C_γ_, Asp85 and Asp212 from C_β_, and internal water molecule 401. This was followed by a harmonic vibrational calculation, restricted to the retinal and Lys216 (from C_γ_). From the resulting 158 vibrational modes, 6 of them had clear corresponding bands in the experimental K-BR spectrum (Fig. 3). Mode 86 is mostly a C10-C11 stretching; mode 89 is mostly a C14-C15 stretching; mode 92 is mostly a C8-C9 stretching; and mode 93 as mostly a 12C-13C /14C-15C stretching (Fig. 3E). In addition, we identified mode 72 as a 19C+20C methyl rock and mode 82 as a H15C=NH HOOP (Fig. 3E).

For the anharmonic vibrational calculations, we computed potential energy surfaces (PES) and dipole moment surfaces (DMS) along some selected normal mode coordinates. In scheme A (Fig. 3A,B), 116 coordinates with higher energy were set to be active. Anharmonic frequencies and intensities were calculated by vibrational self-consistent field (VSCF) with 2nd-order quasi-degenerate perturbation theory (VQDPT2), based on PES and DMS derived at the level of B3LYP-D3/cc-pVDZ. In the scheme B (Fig. 3C,D), we used a more accurate basis sets to calculate PES and DMS, B3LYP-D3/cc-pVTZ, although limited to only 8 selected coordinates: those presented in Fig. 3E plus mode 70 (6C-12C HOOP) and mode 136 (N-H stretch). In addition, anharmonic frequencies and intensities were obtained by VSCF with vibrational configuration interaction (VCI), more accurate but computationally more expensive than VSCF + VQDPT2.

Although the main purpose of these two anharmonic calculations was to assist us in the assignment of experimental bands to overtone and combination transitions, we first compared their performance in reproducing assignments of bands to fundamental vibrational transitions of all-trans PSB retinal in BR. To improve the comparison between experimental and calculated spectra, we took into consideration that the experimental data for the BR state comes from a K-BR difference spectrum. Consequently, bands unchanged between the BR and K states are not experimentally observed, even if present in the BR state. One example are vibrations localized in the β-ionone ring of the retinal. For that reason, we also reconstructed anharmonic IR spectra selecting only retinal vibrational modes that we experimentally know to change between the BR and K states (Fig. 3A-D, red vertical lines and blue traces).

Scheme A predicts vibrational modes 86 (10C-11C), 89 (14C-15C) and 93 (12C-13C/14C-15C) at 1211, 1232 cm^−1^ and at 1269 cm^−1^, respectively (Fig. 3A). The agreement with experimental bands from 10C-11C, 14C-15C, and 12C-13C/14C-15C stretches, at 1169, 1202 and 1255 cm^−1^, is good, with an error of +14-30 cm^−1^ (Fig. 3A). In addition, the relative areas of these bands (1 / 2.0 / 1.2) are well reproduced in the calculation (1 / 2.3 / 1.7), as collected in Table 1. The main failure of this calculation is that it assigns a tiny relative intensity to mode 92 (Fig. 3A), around 10 times smaller than experimentally observed for the C8-C9 stretch (see Table 1). Furthermore, it predicts the frequency of this mode to lay between those of modes 89 and 86, instead of laying between those of modes 93 and 89, as expected from the experimental data. Regarding modes 72 and 82, their fundamental is predicted at 1002 and at 1040 cm^−1^, respectively (Fig. 3A), very close to the experimental values for the H15C=NH HOOP (1008 cm^−1^) and the 19C/20C methyl rock (1050 cm^−1^). In summary, calculation of scheme A reproduced well bands from fundamental vibrations of all-trans PSB retinal in the BR state, except for the 8C-9C stretching (see Fig. 3A and Table 1).

The commented limitation of scheme A led us to scheme B, more accurate but with a reduced dimensionality to compensate for the additional computational effort. The agreement between the experimental frequencies of retinal C-C stretches and those calculated for modes 86 (10C-11C), 89 (14C-15C) and 93 (12C-13C /14C-15C) was good, even though the error made (+42-50 cm^−1^) was clearly larger than for scheme A (compare Fig. 3A and Fig. 3C). The calculated relative intensity of these modes (1 / 0.76 / 0.86), although reasonable, was also worse than for scheme A (Table 1). Regarding modes 72 and 82, this calculation provided frequencies at 1034 cm^−1^ and at 1108 cm^−1^ (Fig. 3C), off by +26 cm^−1^ and +58 cm^−1^. Finally, regarding mode 92 (8C-9C), problematic in scheme A, its frequency was correctly predicted to lay between modes 93 and 89, with an absolute error of +47 cm^−1^, consistent with the ∼+50 cm^−1^ error made for the rest of C-C modes (Fig. 3C). Still, the calculated relative intensity of mode 92 was 3.5 times higher than expected from the experimental data (Fig. 3C and Table 1).

Overall, both anharmonic calculations successfully reproduced frequencies and intensities of experimental bands from fundamental transitions of all-trans PSB retinal in the BR state. Among them, the performance of scheme A was somehow better, but it clearly failed for mode 92 (8C-9C), for which scheme B, with the use of larger basis sets, provided far more reasonable parameters. This indicates that for overtone and combination transitions involving mode 92, the predictions from scheme B are likely to be more reliable.

### 3.5. Assignment of overtones and combinations of PSB retinal C-C stretches

In the 2550-2325 cm^−1^ region of the K-BR spectrum (Fig. 2, red trace), we detect four negative bands (2336, 2402, 2452 and 2504 cm^−1^) and two positive bands (2385 and 2365 cm^−1^). In addition, two additional negative bands are resolved at 2427 and 2416 cm^−1^ after band-narrowing (Fig. 2, pale red trace). A tiny negative band is also observed at around 2478 cm^−1^. We have tentatively assigned all these bands to overtones of C-C stretches, except for bands at 2478 cm^−1^, 2452 cm^−1^ and 2416 cm^−1^, assigned to combinations of C-C stretches (see Fig. 2 and Table 2).

To validate and to complement these assignments, or correct them if needed, we performed two anharmonic calculations, introduced in the previous section. Figure 3B,D compares experimental and calculated spectra for all-trans PSB retinal in the 2650-2200 cm^−1^ region, containing overtone and combination modes. As explained in detail in the previous section, our anharmonic calculations overestimated fundamental νC-C transitions by ∼15-30 cm^−1^ (scheme A) or by ∼40-50 cm^−1^ (scheme B) and, thus, frequencies of overtones and combination bands are expected to be overestimated by 30-60 cm^−1^ (scheme A) or by 80-100 cm^−1^ (scheme B).

From the seven experimental negative bands, four of them are well reproduced in the calculation from scheme A (Fig. 3B and Table 2). Their frequencies are 31-60 cm^−1^ upshifted in the calculation, as expected. The experimental band at 2336 cm^−1^, predicted at 2395 cm^−1^, originates from the overtone of mode 86 (10C-11C). The experimental band at 2402 cm^−1^, predicted at 2462 cm^−1^, can be assigned to the combination of mode 92 (8C-9C) and mode 89 (14C-15C), overlapping with the overtone of mode 89 (14C-15C), the latter less intense. We attribute the experimental band at 2504 cm^−1^, predicted at 2535 cm^−1^, to the overtone of mode 93 (12C-13C / 14C-15C). Finally, the experimental band at 2452 cm^−1^, predicted at 2498 cm^−1^, can be assigned to a combination of mode 93 (12C-13C / 14C-15C) and mode 89 (14C-15C).

The calculation from scheme A also displayed some apparent inconsistences with the experimental data. It predicts a rather intense transition at 2430 cm^−1^, originating from the combination of mode 89 (14C-15C) and mode 86 (10C-11C) (Fig. 3B). However, the corresponding experimental band, expected at ∼2390-2370 cm^−1^, is not observed. While this could be due to an error in the calculation, we cannot completely discard that this band is present but cancelled by the more intense positive bands at 2385 and 2365 cm^−1^ from 13-cis PSB retinal in the K state (Fig. 3B). A more serious issue comes by the failure of scheme A in reproducing three experimental negative bands: 2427 cm^−1^, tentatively assigned to the overtone of the fundamental at 1216 cm^−1^ (8C-9C), and at 2478 cm^−1^ and at 2416 cm^−1^ (Fig. 2, red spectrum), these two bands assigned to the combination of the fundamental at 1216 cm^−1^ (8C-9C) with either the fundamental at 1255 cm^−1^ (12C-13C / 14C-15C) or with the fundamental at 1201 cm^−1^ (14C-15C), respectively (see Table 2). Actually, the calculation from scheme A predicted a negligible intensity for the overtone of mode 92 (8C-9C), as well as for the combination of mode 92 with mode 89 (14C-15C) and of mode 92 with mode 93 (12C-13C / 14C-15C) (Fig. 3B and Table 2). It also wrongly predicted the frequency of the overtone of mode 92 at 2450 cm^−1^, between the overtones of modes 89 and 86 (see Fig. 3B). In summary, the calculation from scheme A had problems reproducing the overtone of mode 92, as well as several of its combinations with other modes.

Scheme B reproduced all seven experimental negative bands in the 2550-2325 cm^−1^ region (Fig. 3D). The predicted bands appear +86-106 cm^−1^ upshifted in the calculation (Fig. 3D), fully consistent with the upshift of fundamental vibrations by 42-51 cm^−1^ (Fig. 3C) In agreement with scheme A, the experimental bands at 2335, 2402 and 2593 cm^−1^ can be assigned to the overtones of mode 86 (10C-11C), mode 89 (14C-15C), and mode 93 (12C-13C / 14C-15C), respectively (Table 2), and the experimental band at 2452 cm^−1^ to the combination of mode 93 (12C-13C / 14C-15C) and mode 89 (14C-15C). Regarding bands not explained by the previous calculation, the weak experimental band ∼2427 cm^−1^, predicted here at 2529 cm^−1^ can be assigned to the overtone of mode 92 (8C-9C). The nearby experimental band at 2416 cm^−1^, with a predicted frequency at 2519 cm^−1^ (not marked in Fig. 3D by lack of space but included in Fig. 2 and Table 2), can be explained by the combination of mode 92 (8C-9C) and mode 89 (14C-15C). Finally, the tinny experimental band at 2478 cm^−1^, is consistent with the combination of mode 93 and mode 92, with a predicted frequency at 2567 cm^−1^ (Table 2).

Scheme B predicts two intense vibrational transitions with no apparent experimental counterpart bands. One is, like in scheme A, the combination of mode 89 and mode 86, predicted at 2466 cm^−1^ (Fig. 3D and Table 2). Given the systematic error in this calculation, the corresponding experimental negative band would be expected at ∼2366-2380 cm^−1^. The other transition is the combination of mode 92 and mode 89, at 2478 cm^−1^ (Fig. 3D and Table 2), with its experimental negative counterpart band expected at ∼2378-2392 cm^−1^. It is possible that these two combination bands are actually present but not observed in the K-BR spectrum because they haven an intensity than calculated, being fully cancelled by overlapping positive bands from the K intermediate. Transitions involving mode 92 weaker than calculated are reasonable when considering how scheme B overestimated the relative intensity of its fundamental.

In summary, our anharmonic calculations support that four of the seven negative experimental band between 2550-2325 cm^−1^ are overtone vibrations of C-C stretches from all-trans PSB retinal (Table 2). Both calculations agree that the band at 2335 cm^−1^ corresponds to the overtone of the fundamental at 1169 cm^−1^ (10C-11C); that the band at 2402 cm^−1^ corresponds to the overtone of the fundamental at 1202 cm^−1^ (14C-15C); and that the band at 2504 cm^−1^ corresponds to the overtone of the fundamental at 1255 cm^−1^ (12C-13C/14C-15C). The calculation from scheme B indicates that the band at 2427 cm^−1^ corresponds to the overtone of the fundamental at 1217 cm^−1^ (8C-9C). As for combination bands, the negative band at 2453 cm^− 1^ can be assigned to the combination of the fundamentals at 1255 cm^−1^ (12C-13C / 14C-15C) and at 1202 cm^−1^ (14C-15C), accordingly to both calculations. The band at 2416 cm^−1^ corresponds to the combination of the fundamentals at 1216 cm^−1^ (8C-9C) and 1201 cm^−1^ (14C-15C), and the band at 2478 cm^−1^ to the combination of the fundamentals at 1255 cm^−1^ and at 1216 cm^−1^, accordingly to the calculation from scheme B. These assignments are collected in Table 2.

Regarding potential overtones bands from C-C stretches of 13-cis PSB retinal, we observe a positive band at ∼2385 cm^−1^ and at ∼2365 cm^−1^ in the K-BR spectrum (Fig. 2, red trace). These bands are akin to the positive band at ∼1194 cm^−1^ and the shoulder at ∼1182 cm^−1^, but at roughly twice their wavenumber. Therefore, we assign them to overtones from 14C-15C and 10C-11C stretches of 13-cis PSB retinal (Fig. 2, blue).

### 3.6. Anharmonic and coupling mechanical constants for PSB retinal C-C stretches

The mechanical anharmonic constant for a vibrational mode can be determined as the fundamental minus half the first overtone frequency. On the other hand, the mechanical coupling constant between two vibrational modes is given by the sum of their fundamentals minus their combination frequency. To visualize these constants, we overlayed the 1270-1150 cm^−1^ and 2540-2300 cm^−1^ regions of the K-BR difference spectrum, before (Fig. 4A) and after (Fig. 4B) band-narrowing. To quantify more accurately the anharmonic constants, we determined the frequency of fundamental, combination and overtone bands from band-narrowed K-BR spectra (shown in Fig. 4B and collected in Table 2 for the negative bands).

**Figure 4.**
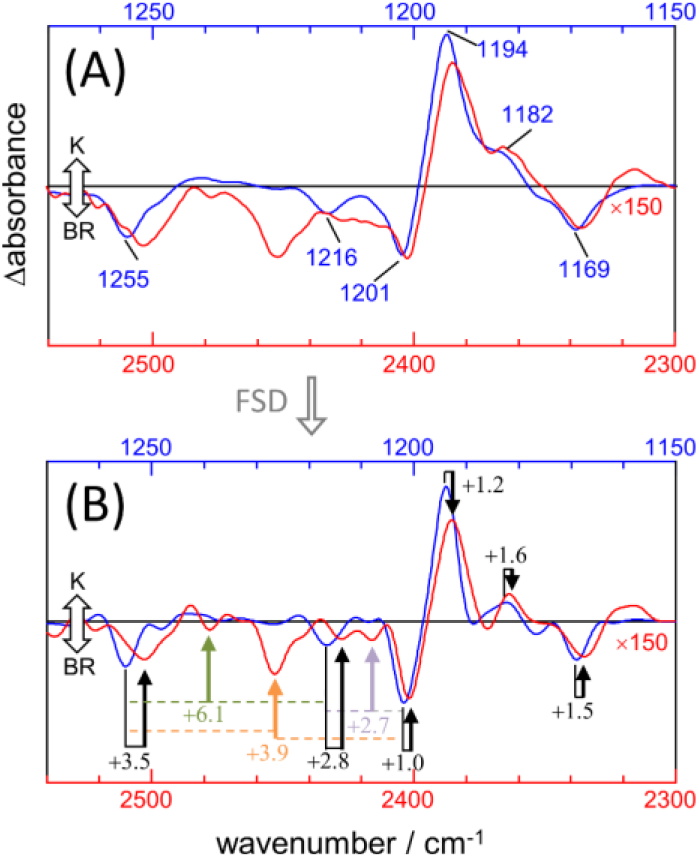
Comparison between νC-C fundamentals (blue traces) and νC-C overtones (red traces) in the K-BR spectrum. (a) Experimental K-BR difference FT-IR spectrum in the 1275-1150 cm^−1^ (blue) and 2550-2300 cm^−1^ (red) region. Wavenumbers of bands from fundamental transitions are indicated (see Table 1 for their assignments). (b) Band-narrowed K-BR spectrum using a narrowing factor of 1.6 and a Lorentzian width of 7 cm^−1^ for the 1275-1150 cm^−1^ region and of 14 cm^−1^ for the 2550-2300 cm^−1^ region. Overtone bands are labelled with their estimated mechanical anharmonic constant (black) and combination bands with their mechanical coupling constant (color) constants (see Table 2 for their assignments). Note that in both (a) and (b) the overtone region is scaled by 150.

For the 14C-15C and 10C-11C vibrations of all-trans PSB retinal, with fundamentals at ∼1201 and ∼1169 cm^−1^ (Fig. 4A), respectively, the mechanical anharmonic constants are +1.0 cm^−1^ and +1.5 cm^−1^ (Fig. 4B). For 13-cis PSB retinal in the K intermediate the coupled 14C-15C and 10C-11C vibrations, with fundamentals at 1194 and 1182 cm^−1^, display mechanical anharmonic constants of 1.2 cm^−1^ and 1.6 cm^−1^, respectively (Fig. 4A, right). Thus, the mechanical anharmonic constant for the 14C-15C and 10C-11C stretches are quite low, regardless of the isomerization state of the PSB retinal: 1.0-1.5 cm^−1^ in the BR state (all-trans), and 1.2-1.6 cm^−1^ in K intermediate (13-cis).

Returning to all-trans PSB retinal, the vibrational mode containing both 12C-13C and 14C-15C contributions, with the fundamental at ∼1255 cm^−1^ (Fig. 4A) displays a mechanical anharmonic constant of +3.5 cm^−1^ (Fig. 4B). For the C8-C9 stretching, contributing to the fundamental at 1216 cm^−1^, its mechanical anharmonic constant is +2.8 cm^−1^ (Fig. 4B). Regarding mechanical coupling constants, the one between the fundamental at 1201 cm^−1^ (14C-15C) and at 1255 cm^−1^ (12C-13C / 14C-15C) is of +3.9 cm^−1^ (Fig.4B); between the fundamental at 1201 cm^−1^ (14C-15C) and at 1216 cm^−1^ (8C-9C) of +2.7 cm^−1^; and between the fundamental at 1255 cm^−1^ (12C-13C / 14C-15C) and at 1216 cm^−1^ (8C-9C) of +6.1 cm^−1^.

The mechanical anharmonic and coupling constants for all-trans PSB retinal determined from the scheme A are in reasonable agreement with those determined experimentally (Table 2). In particularly, the anharmonic constant for the of mode 86 (10C-11C) was calculated to be +0.9 cm^−1^, comparing well with the experimental value +1.5 cm^−1^. For other transitions the error was somehow larger: for mode 89 (14C-15C) and for mode 93 (12C-13C / 14C-15C) the mechanical anharmonic constant was calculated to be +3.9 cm^−1^ and +1.1. cm^−1^, but experimentally determined to be of +1.0 cm^−1^ and +3.5 cm^−1^, respectively (Table 2). The coupling constant for the combination of modes 89 (14C-15C) and 93 (12C-13C / 14C-15C) was calculated to be +2.1 cm^−1^, not far from its experimental value of +3.9 cm^−1^ (Table 2).

Regarding mechanical anharmonic constants predicted by scheme B, they were also very small, like the experimental ones, but negative, ranging from –1.8 to –0.1 cm^−1^ (Table 2). The mechanical coupling constants were, likewise, predicted to be small but negative, ranging from – 4.4 to –3.1 cm^−1^ (Table 2).

### 3.7. Width and relative intensity of bands from overtone transitions of C-C stretches

Figure 4a compares the fundamental and overtone/combination region of C-C stretches of the retinal. These two regions were scaled to make the band at 1169 cm^−1^ (10C-11C) and its overtone at 2336 cm^−1^ to be of roughly the same intensity. Thus, it can be visually appreciated that the overtones of retinal C-C stretches are roughly 150 times weaker in intensity that their fundamentals. More relevant are area ratios between fundamental and overtones, being 60, 110, 80, 85, 70 and 50 for the fundamental vibrations at 1255 (–), 1216 (–), 1201 (–), 1194 (+), 1182 (+) and 1169 (–) cm^−1^, respectively (Table 2). In contrast, overtones were calculated to be between 250 to 1800 weaker than their corresponding fundamental transitions in scheme A, and between 240 to 1700 in scheme B (Table 2). Thus, C-C overtones are ∼4-20 more intense in the experiments than in our anharmonic calculations, a notable difference that calls for an explanation.

Figure 4a also informs us that overtone bands from C-C vibrations of the PSB retinal are roughly twice broader than fundamentals bands, except for the overtone of the 1255 cm^−1^ fundamental, which appears 3-4 times broader. Thus, although the separation of bands is twice larger for overtones than for fundamental vibrations, this is of no help to improve their band resolution, particularly when considering that overtones bands co-exist in this region with combination bands.

### 3.8. Combination bands of retinal C-C stretches with CH_3_ rocks and HOOPs

The negative band at 1008 cm^−1^ corresponds to the symmetric in-plane rocking combination of 19C and 20C methyl groups from the BR state (Smith et al., 1987; Yamazaki et al., 1995), akin to mode 72 in our calculations (Table 1). Scheme A predicts the combination of mode 72 with the three more intense C-C modes of all-trans PSB retinal (modes 93, 89 and 86), giving rise to combination bands at 2270, 2235, and 2202 cm^−1^ (Table 2). This assignment agrees with the experimental negative bands at 2257 and 2212 and 2176 cm^−1^, which can be explained by the combination of the fundamental at 1008 cm^−1^ with fundamental transitions from C-C stretches at 1255, 1202 and 1169 cm^−1^, respectively. According to the above assignment, the experimental mechanical coupling constants between the methyl rock and these C-C stretches are +6.2, –1.8 and +1.4 cm^−1^, respectively. The prediction from scheme B is similar, except for the fact that mode 72 is predicted to couple with mode 92 instead of with mode 89. Accepting this last assignment, the experimental band at 2212 cm^−1^ would result from the combination of the fundamentals at 1008 cm^−1^ and at 1216 cm^−1^, with a mechanical coupling constant of +13 cm^−1^. Between these two possibilities, we think that in this case it is more reasonable to assignment the band at 2212 cm^−1^ to the combination of the fundamentals at 1008 cm^−1^ and 1201 cm^−1^ (modes 72 and 89), as suggested by scheme A.

Positive bands at 2165, 2149 and 2135 cm^−1^ fit well with the combination of the 14C-15C + 10C-11C stretch at 1194 cm^−1^ with in-phase HOOP bands at 974, 956 and 941 cm^−1^ (Fig. 2). In spite the high relative intensity of these combination bands, the derived mechanical coupling constants are very small, between –0.2 and +1.0 cm^−1^.

### 3.9. Unassigned bands between 2100 and 1850 cm^−1^

A broad negative and a positive band are resolved at around 2070 (–) cm^−1^ and around at 2000 (+) cm^−1^, respectively, with a fine structure revealed after band-narrowing by FSD (Fig. 2, bottom). Given their frequency, they might arise from combination bands between out-of-phase HOOP bands, located between 850-750 cm^−1^ (Smith et al., 1987), and C-C stretches. Relatively narrow alternating positive and negative bands are observed between 1950 and 1850 cm^−1^. Although from their frequency they could be overtones of in-phase HOOP bands, they do not display any clear resemblance to HOOPs bands from fundamental transitions. Alternatively, these bands could also originate from the coupling of C-C stretching vibrations with some fundamental vibrations at around 700 cm^−1^. Experiments with isotope labelled retinal will be needed for the assignment of bands in this spectral region to specific overtones/combination vibrational transitions.

## 4 General Discussion

Bands in the mid-IR region of proteins are, by default, assumed to corresponds to fundamental transitions from molecular vibrations. While this assumption is often well justified, in this work we have shown that the light-induced FT-IR difference spectrum of bacteriorhodopsin contains bands in the mid-IR region coming not from fundamental but from overtones and combination transitions. In particular, we have shown that bands between 2540 and 2300 cm^−1^ originate from overtones and combination bands from retinal C-C vibrations, and bands between 2300 and 2100 cm^−1^ result from transitions involving the combination of retinal C-C vibrations with either methyl rocks or hydrogen-out-of-plane vibrations.

Another notable observation of the present work is the fact that retinal C-C stretching modes are, apparently, close to be mechanically harmonic, with mechanical anharmonic constants for the 14C-15C and 10C-11C stretches between 1.0 and 1.6 cm^−1^ for both all-trans PSB and 13-cis PSB. Despite these small values, the overtones from 14C-15C and 10C-11C stretches display areas only ∼50-85 smaller than their corresponding fundamentals. To have a reference, the anharmonic constant for CO bound to myoglobin is 12-14 cm^−1^, almost 10 times larger than for the retinal 14C-15C and 10C-11C stretches, but the area of its overtone is ∼100 smaller than for its fundamental (Sage et al., 2011). To give an explanation, we have to note that the intensity of overtone bands depends both on the mechanical anharmonic constant (through the wavefunction overlap of the fundamental and the second excited vibrational state) and on the electrical anharmonicity (related to δμ^2^/δ^2^Q, where Q is the mode coordinate and μ is the molecular dipole moment) (Sandorfy et al., 2006). Thus, it is reasonable to conclude that, given the low mechanically anharmonicity of retinal C-C stretches, leading to close to harmonic wavefunctions, the rather high intensity of the overtone bands can only be explained if the electrical anharmonicity of the retinal C-C stretches is unusually large.

We asked ourselves how similar are mechanical anharmonic constants and fundamental/overtone area ratios for C-C stretches in the PSB retinal of BR when compared to C-C stretches from other molecules. This question is difficult to answer though. The reason is that C-C stretches have a very low associated transition dipole moment and, thus, very weak intensities for fundamental (and overtone) transitions. Indeed, we have not been able to find examples in the literature for experimental mechanical anharmonic constants of C-C stretches or fundamental/overtone ratios. The case of PSB retinal is peculiar. Because of the pi-conjugated system, when the SB is protonated the positive charge is delocalized along the polyene chain, with a gradient: the local charge is highest at the N atom, decreasing in the direction to the ring (Babitzki et al., 2009). As a result, the C-C (and C=C) bonds are polarized and their stretching vibrations are associated with a relatively large change in the dipole moment, giving IR intensity to the fundamental transitions (Babitzki et al., 2009). In agreement, the experimental IR intensity of the C-C stretching vibrations of the PSB retinal drops dramatically when deprotonated, as in the M intermediate of BR (Maeda, 1995).

We believe that the charge delocalization in the PSB retinal explains not only the large transition dipole moment of the C-C stretching vibrations (i.e., explain the IR intensity for the fundamental transitions), but explains as well the electrical anharmonicity of the C-C stretches (i.e., the overtone intensity). If this interpretation is correct, overtone and combination bands from C-C stretches will be sensitive to the charge distribution of the PSB retinal, which in turn is sensitive to local electric fields (Babitzki et al., 2009), and thus, to the environment and nature of the binding pocket of the PSB retinal. In this respect, it is interesting to note that the mechanical and electrical anharmonicity of retinal C-C stretches is similar but not identical after retinal isomerization, as deduced from the similar but not identical anharmonic constants and fundamental/overtone areas for positive and negative bands in the K-BR FT-IR difference spectrum. Further work should focus in resolving overtone vibrations for the rest of the intermediates in the photocycle of BR, providing a more ample view on the subject. We should also note that the present results have been obtained at 80 K. Another pending question is if the wavenumber and intensity of the overtone and combination bands from the BR state and, thus, the mechanical and electronic anharmonicities of the vibrations of the PSB all-trans retinal, might be affected by the temperature at which the photocycle is initiated. If so, this could indicate temperature-dependent changes in the retinal binding pocket. A final question that requires further investigation is the role of the environment on the mechanical and electronic anharmonicities of the PSB retinal. In this respect it would be very interesting to resolve and characterize overtone and combination bands of PSB all-trans retinal in solution. In the past we have studied a model for PSB retinal in solution by UV-vis spectroscopy (Lórenz-Fonfría et al., 2010), but so far we have not succeeded in conducting similar studies by FT-IR spectroscopy.

The present results also provide information for mechanical coupling constant between vibrational modes. Coupling constants between C-C stretches are between +2.7 and +6.1 cm^−1^. Coupling constants between C-C stretches and C-CH_3_ (methyl rocks) or C-H wags (HOOP vibrations) of the retinal are generally smaller, in most cases in the order or lower than 2 cm^−1^. The observation of intense combination bands between the C-C and HOOP vibrations of the retinal in the K intermediate, in spite their coupling constant being smaller than 1 cm^−1^, indicate that both vibrations are electronically coupled. In other words, δμ^2^/δQ_*i*_Q_*j*_ ≠ 0, meaning that oscillations along the C-C coordinate affect how the dipole moment changes along the HOOP vibration coordinate, and vice versa, resulting in their coupling. An interesting question is whether the electronic coupling reported between C-C and HOOP vibrations could allow to transfer energy from one vibrational mode to the other, and if this transfer could help or play a role in the thermal relaxation of the retinal following its photoisomerization from all-trans to 13-cis.

The in-phase C=C stretching vibration of PSB retinal in the BR, at 1526 cm^−1^, has been shown by 2D-IR spectroscopy to have a mechanical anharmonic constant of 7 cm^−1^ (Andresen and Hamm, 2009), substantially larger than the values we determined here for the C-C stretches. Why are the C-C stretches mechanically more harmonic than the C=C stretches? One difference between both types of vibrations is that the modes assigned to C-C stretches also involve in-plane bending models of the hydrogen atoms attached to the polyene chain (CCH in-plane rocks), as revealed by normal mode calculations and supported by ^13^C and ^2^H labelling (Smith et al., 1987; Babitzki et al., 2009). Thus, it is possible that the relatively low mechanical anharmonicity of the C-C stretches could be, at least partially, caused by the contributions from these bending modes to the vibrational modes, which are expected to behave more harmonically than stretching vibrations. A way to test this hypothesis in the future could be experiments with perdeuterated retinal, uncoupling the C-C stretches from in-plane bending modes.

## Author Contributions

VALF conducted experiments and wrote a draft of the manuscript, KY performed calculations, SI conducted experiments, HK provided sample and access to equipment, all authors discussed the results and edited the final version of the manuscript.

## Data Availability Statement

Data included in the figures of the manuscript (measured, analyzed, or calculated) can be found in Mendeley Data [DOI: 10.17632/n5t4jh5cm9.1].

