## Supplemental Figures for "Retinal vibrations in bacteriorhodopsin are mechanically harmonic but electronically anharmonic: evidence from overtone and combination bands"

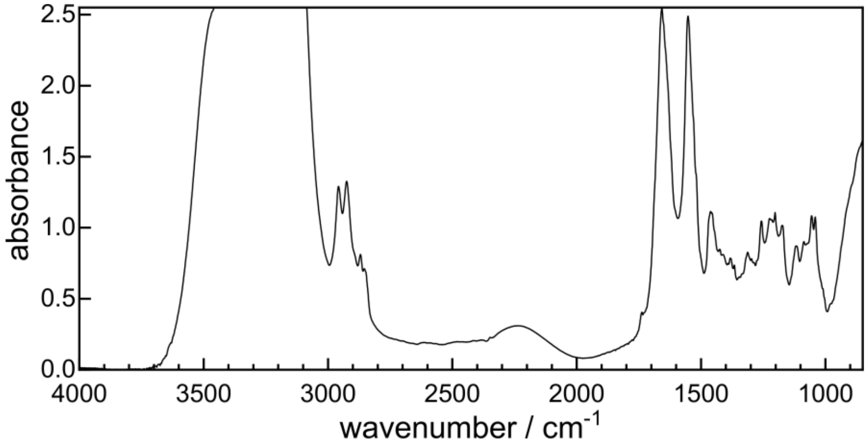


**Supplementary Figure 1.** Absorption FT-IR spectrum at 80 K of the hydrated film of BR in purple membrane used in the present study. The film was thicker than usually, leading to an absorbance above 2 in several spectral regions, but also to increase of the signal from overtone and combination bands.

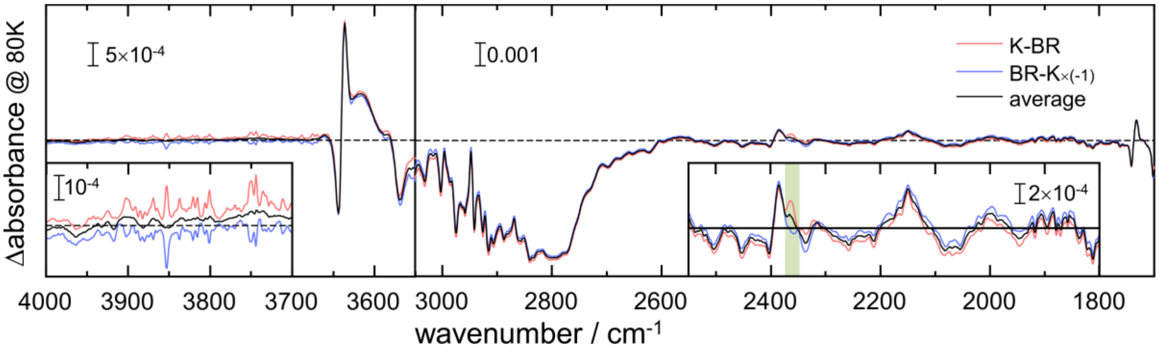


**Supplementary Figure 2.** K-BR (red trace), minus BR-K (blue trace), and the averaged K-BR FT-IR difference spectrum (black trace). Note that both K-BR (red trace) and minus BR-K (blue trace) difference spectra have identical signals but opposed vapor and $CO_2$ contributions (see left insert and green area in right insert, respectively), with the average cancelling both contributions.

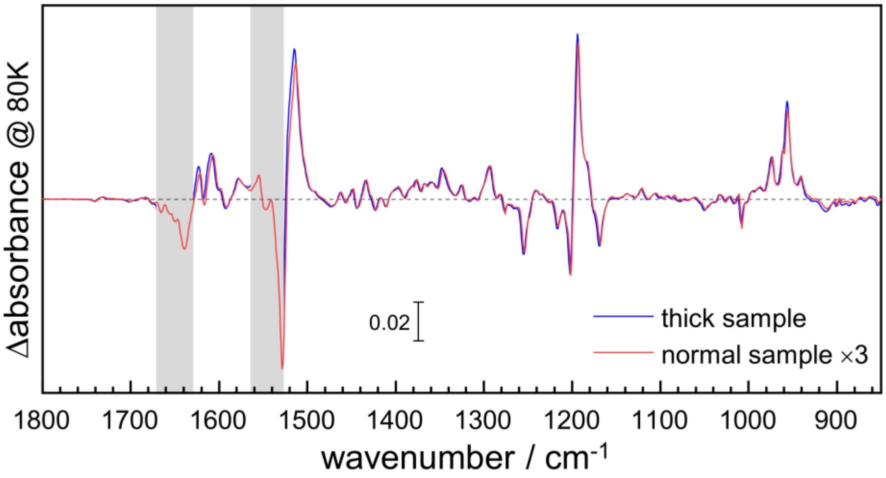


**Supplementary Figure 3.** Light-induced K-BR difference spectrum by FT-IR spectroscopy at 80K, obtained using a normal (red trace) or a thick (blue trace, see Fig. S1) film sample. For the thick sample, the absorption changes in strongly absorbing regions were not reliable (indicated by gray vertical lines). However, it provided three times larger signals (and better signal-to-noise) in other regions, without any significant spectral alterations.

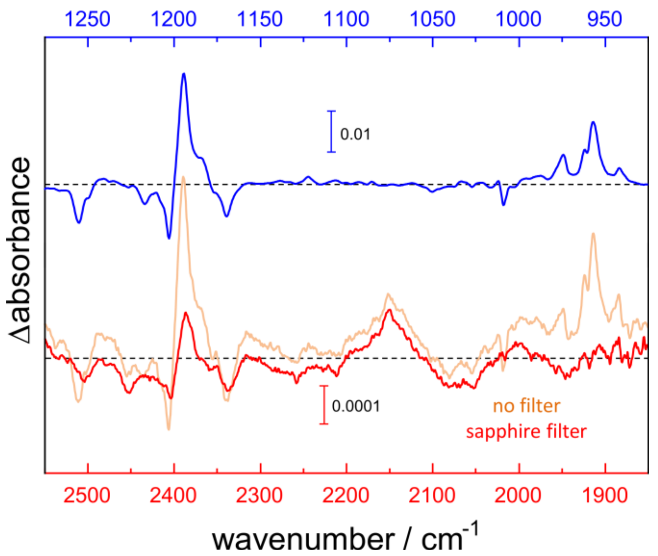


**Supplementary Figure 4.** Comparison of the 2550-1850 $cm^{-1}$ (light red trace) and the 1275-925 $cm^{-1}$ (blue trace) regions of a K-BR difference spectrum obtained with a sample of normal thickness, using an old FT-IR spectrometer with an optical design not optimized to prevent back-reflections to reach the detector. The same spectrum was obtained placing a sapphire window in the IR beam path (red trace), which removes any double-modulation artefactual bands from around 3200 to 1800 $cm^{-1}$.

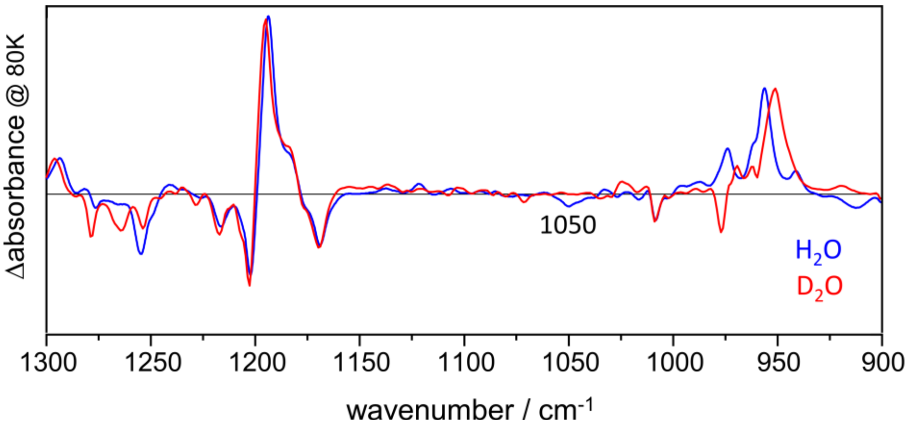


**Supplementary Figure 5.** K-BR difference FTIR spectrum, obtained with a sample hydrated with $H_2O$ (blue) or $D_2O$ (red). Hydration with $D_2O$ only deuterates the N-H group from the retinal. The disappearance of the negative band at 1050 $cm^{-1}$ in $D_2O$ indicates, also given its wavenumber, that this band originates from a vibration with contributions from a N-H bend.
